# Mild subcortical stroke induces widespread astrogliosis independent of microglia and age

**DOI:** 10.1101/2025.07.18.665576

**Authors:** Teresa L. Stackhouse, Bradley D. Marxmiller, Steve J. Sullivan, Heather L. McConnell, Laura M. Knittel, Rachel De La Torre, Alexandra Houser, Wenri Zhang, Randall L. Woltjer, Anusha Mishra

**Author notes:** **Correspondence:** Anusha Mishra, Ph.D. Department of Neurology Jungers Center for Neurosciences Research 3215 S.W. Pavilion Loop, Portland, OR 97239.

## Abstract

Ischemic stroke induces a plethora of pathophysiological changes, including neuroinflammation and chronic cerebrovascular dysfunction. In humans, even small, silent strokes can trigger these pathologies, which can spread to brain regions far beyond the infarct and persist chronically, ultimately worsening prognosis and increasing the risk for vascular dementia and Alzheimer’s disease. The cause of this extensive pathology is unknown, but reactive astrocytes and microglia are likely contributors. Here, we describe an optimized short-duration middle cerebral artery occlusion model that produces a clinically relevant small stroke confined to subcortical regions, similar to most silent strokes in humans. We termed this model the mild subcortical infarct (MSCI). We then mapped the spatiotemporal extent of reactive astrocytes and microglia during the sub-acute period (1, 3, and 7 days) following MSCI. We observed that reactive astrogliosis develops more rapidly and spreads more extensively, permeating the entire middle cerebral artery territory, compared to the reactive microglial response following this small infarct. Microglial depletion resulted in larger infarct sizes but did not prevent the reactive astrocytes, suggesting that ischemia-driven astrogliosis is largely microglia-independent. Lastly, we show that aging mice exposed to MSCI exhibit a comparably strong response of reactive astrocytes and microglia as young mice. We propose that MSCI is a novel and valuable model for examining the subtle yet highly important chronic effects of stroke. It may be especially useful for investigating the influence of reactive astrogliosis on pathologies like neuroinflammation and cerebrovascular dysfunction in regions distal from the primary injury site.

## 1. INTRODUCTION

Stroke is a leading contributor to major disability in the aging population (Benjamin et al. 2018) and one of the strongest risk factors for dementia (Hachinski 2018), a growing concern for the aging population worldwide. Understanding the complexities of stroke-induced neurological dysfunction is important for discovering new interventions to improve clinical outcomes and quality of life (Feske 2021).

Ischemia and the resulting cell death (infarct) can trigger gliosis, a process wherein astrocytes and microglia proliferate and undergo changes in their morphological and expression profiles. These changes are mainly geared toward the protection of tissue, likely to repair and maintain the infarct border, preventing further damage (Burda and Sofroniew 2014; Amantea et al. 2015). For example, microglia clear dead cells and debris via phagocytosis (Jia et al. 2021), while astrocytes form glial scars that limit the spread of injury (Amantea et al. 2015; Anderson et al. 2016). However, reactive microglia and astrocytes may also release detrimental signals, such as pro-inflammatory cytokines that can exacerbate neuronal damage in the long term (Gong et al. 2023; Amantea et al. 2015). After ischemic infarction, gliosis spreads beyond the immediate infarct and its border regions in an outward gradient, affecting otherwise intact tissue (Gong et al. 2023; Barreto et al. 2011). However, its precise spatiotemporal extent remains unclear. In certain pathological conditions, microglial activation induces specific subtypes of reactive astrocytes, as seen in response to lipopolysaccharide (LPS), a model of bacterial infection (Liddelow et al. 2017), and traumatic brain injury (Witcher et al. 2018). Whether microglia reactivity is necessary for reactive astrogliosis in other pathological contexts, such as after stroke, is also unknown.

Many ischemic stroke studies have focused on the biochemical mechanisms underlying infarct development during the acute period. However, the chronic effects of stroke on areas distant from the initial infarct remain poorly understood, despite long-term deficits extending beyond infarcted regions. Areas distant from the infarct site exhibit abnormalities such as decreased neurovascular coupling (Li et al. 2021), vascular dysfunction, and cognitive deficiencies (Levine et al. 2015; Mijajlovic et al. 2017). Many patients who survive stroke later experience small, silent, and recurring infarcts that can contribute to vascular dementia and Alzheimer’s disease (Miklossy 2003). Notably, subcortical damage without direct cortical injury is also highly clinically relevant, as insults in these regions are more closely associated with dementia (van Rooij et al. 2016; Vermeer, Longstreth, and Koudstaal 2007). Given that both glial cells have a role in modulating cerebral blood flow (Mishra et al. 2024; Bisht et al. 2021), long-term gliosis may contribute to neurovascular dysfunction in regions beyond the infarct, resulting in metabolic stress and the development of dementia in later life by altering the homeostatic functions of astrocytes and microglia (Hickman et al. 2018).

The 60-minute transient middle cerebral artery (MCA) occlusion (MCAo) model is widely used to study ischemic stroke (Longa et al. 1989; Sozmen, Hinman, and Carmichael 2012). This model results in an infarct that spans the subcortical regions, including the striatum, and much of the MCA territory of the cortex (Figure 1B, top). These large strokes are considered malignant strokes in humans and are frequently lethal (Barthels and Das 2020; Carmichael 2005).

**Figure 1.**
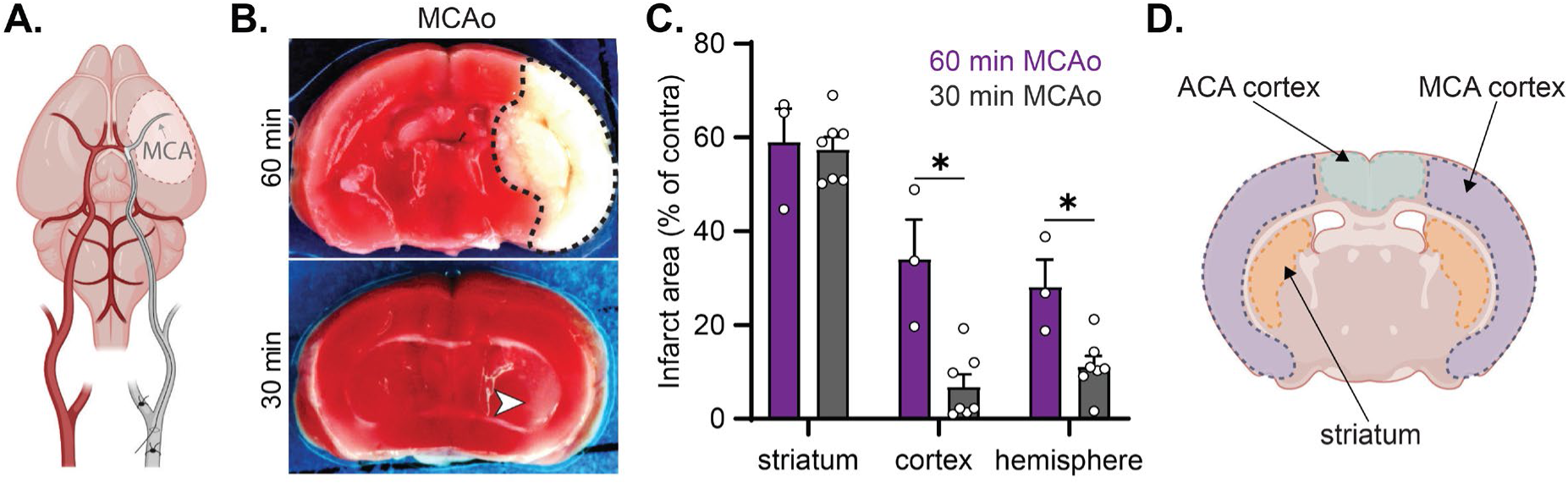
A 30-min middle cerebral artery occlusion produces a mild subcortical infarct. **A.** Schematic of middle cerebral artery occlusion (MCAo) procedure (gray, right) resulting in ischemia within the middle cerebral artery (MCA) territory (dotted outline). **B.** Example brain section stained with 2,3,5-triphenyltetrazolium chloride (TTC) 24 h following a 60-min (top) or 30min (bottom) MCAo. Infarcted tissue does not take up TTC and remains white, revealing a large infarct after 60-min MCAo (top: dotted outline) compared to the diffuse, mild subcortical infarct (MSCI) after 30-min MCAo (bottom: white arrowhead). **C.** Quantification of infarct size 24 h after MCAo. 30-min MCAo induced infarction is similar to that induced by 60-min MCAo in the striatum but much smaller in the cortex, resulting in a smaller overall infarct in the ipsilesional hemisphere. *p<0.05. **D.** Schematic of regions analyzed in Figures. 2–4,7–9, including MCA and anterior cerebral artery (ACA) territories and striatum. Schematics in A and D created with BioRender.com.

Although rodents can survive a 60-minute MCAo for several days, this model is impractical for studying the more chronic effects of stroke. A model that produces a reliable, clinically relevant, subcortical infarct would be valuable for studying how strokes contribute to complex neurodegenerative conditions that develop over long periods, such as vascular dementia and Alzheimer’s disease.

In this study, we describe a new optimized model of stroke for studying the chronic effects of small, mild subcortical infarct (MSCI) using a 30-min MCAo. We observed that MSCI leads to an increase in both reactive astrogliosis and microglial reactivity during the sub-acute period (Birenbaum, Bancroft, and Felsberg 2011), though their spatiotemporal progression patterns differ. Further, we found that while reactive microglia restrict the infarct size, they are not required for inducing reactive astrogliosis after stroke. Additionally, aged mice displayed higher baseline levels of astrocyte and microglia markers, indicating aging-related glial changes, but their response to stroke remained robust and comparable to that of mid-life adults.

## 2. MATERIALS AND METHODS

### 2.1 Animals

All experiments were done in accordance with the policies of Oregon Health & Science University’s Institutional Animal Care and Use Committee. Male and female 4–6 months old C57/B6 mice were used as mid-life adult animals for all experiments, except for the aging experiment (Figures 7–9), where only 20-month-old female mice were available.

### 2.2 Transient MCA occlusion (MCAo)

MCA occlusion was performed following previously reported methods (Longa et al. 1989; Li et al. 2021). For Figure 1, both 60-min and 30-min occlusions were performed, while the remaining study used only 30-min occlusions or sham surgeries. Prior to the occlusion, a laser Doppler flowmeter probe was affixed over the right parietal bone overlying the MCA territory to monitor changes in cerebral blood flow. A ventral midline incision was made over the neck, the right common carotid artery (CCA) bifurcation was exposed by gentle dissection and tissue retraction, and the external carotid artery (ECA) was permanently ligated distal to the occipital artery using electrocautery. The right ECA and internal carotid artery (ICA) were temporarily closed with reversible slip knots before an arteriotomy was made in the ECA stump. A 5.0 nylon (silicone-coated) monofilament (Doccol Corporation) appropriate for the weight of the mouse was inserted into the ICA via the arteriotomy and gently advanced to the ICA/MCA bifurcation to occlude cerebral blood flow to the MCA territory, confirmed by a drop in flow measured by laser Doppler flowmetry (Moor Instruments). After a 30– or 60-min occlusion, the filament was gently retracted to allow reperfusion, the ECA permanently ligated, the slip knot on the CCA removed, and the incision sites sutured closed. The sham surgery involved exposure of the CCA and ECA without any arteriotomy or ligations. Mice remained under anesthesia for the entire 30 min and then the incision site was closed. Sham and MCAo mice were allowed to recover for up to 7 days as indicated by the experimental design.

Infarct size was measured 24 h after MCAo in 2 mm thick coronal brain sections (five total) using 2,3,5-triphenyltetrazolium chloride (TTC) staining and digital image analysis.

Sections were incubated in 1.2% TTC in saline for 15 min at 37°C and then fixed in formalin for 24 h.

### 2.3 PLX-3397 administration

All specialty feed was produced by ResearchDiets. Animals were fed ad libitum and monitored daily for consumption. Open Standard Diet from ResearchDiets (Cat. No. D11112201) was impregnated with Pexidartinib (also called PLX3397, Cat. No. C-1271, Chemgood) at a concentration of 290 mg/kg (Elmore et al. 2014). The control diet was a nutrition-matched Open Standard Diet. Separate cohorts of mice were fed the PLX-3397 impregnated or matching control diet for 3 weeks prior to stroke induction and remained on the same diet until euthanasia and tissue collection.

### 2.4 Perfusion

Mice were transcardially perfused with 1% heparinized saline via a needle placed in the left ventricle, followed by fresh-made room temperature 4% paraformaldehyde (PFA) prepared in phosphate-buffered saline (PBS). Following complete fixation (confirmed by drainage of blood from the cut right atrium, visualization of liver paling, and stiffness of the body), brains were extracted and placed in 4% PFA overnight at 4°C before storing in PBS for subsequent immunohistochemistry analysis.

### 2.5 Immunohistochemistry

Perfusion-fixed brains were paraffin-embedded, 6-µm sections were cut, antigen retrieval was performed using standard methods (30 min incubation in citrate buffer, pH 6.0, at 80°C), tissue sections were blocked with 3% nonfat dry milk in PBS and immunolabeled with primary antibodies: rabbit anti-GFAP, 1:500 (Cat. No. 16825-1-AP, Proteintech); rabbit anti-vimentin 1:500 (Cat No. 10366-1-AP, Proteintech); rabbit anti-Iba1, 1:5000 (Cat. No. 10904-1-AP, Proteintech). Appropriate secondary antibodies conjugated to horseradish peroxidase were then used and reacted with 3,3’-diaminobenzidine (DAB) for visualization. Cell nuclei were lightly counterstained with hematoxylin.

### 2.6 AxioScan Imaging

Immunolabeled slides containing whole brain sections were imaged on a Zeiss Axioscan.Z1 with either a 10x/0.45NA objective and a Hitachi HV-F202 camera (resulting in a pixel size of 0.441 µm x 0.441 µm) or a 20x/0.8NA objective under the same system (resulting in a pixel size of 0.220 µm x 0.220 µm).

### 2.7 Region of interest and image selection

GFAP and vimentin: All analysis was conducted in three regions: 1) the anterior cerebral artery territory of the cortex (termed ACA; a region that does not receive any ischemia), 2) the middle cerebral artery territory of the cortex (termed MCA; a region that experiences transient ischemia), and 3) the intact, non-infarcted portions of the striatum (a region that experiences transient ischemia and borders the infarct). ROIs were selected while avoiding areas with visible MCA infarction, which was identified by loss of H&E labeling and visible cell death. GFAP-positive astrocytes also die within the infarct, providing additional confirmation.

Iba1: Four regions in the ipsilesional hemisphere were selected: 1) the ACA and 2) the MCA, as previously described, 3) the intact, non-infarcted portion of the striatum (termed border striatum), and 4) the infarcted region of the striatum (termed infarct). The infarct was identifiable by high Iba1 signal intensity but was defined as a region without GFAP-labeled cells in the adjacent section (as astrocytes also die within the infarct) to avoid subjective bias.

Regions were identified based on structural morphology using the Allen Brain Reference Atlas. All reasonable effort was made to match coronal sections between animals and across labels. For each region, three representative regions of interest (ROIs; 221^2^ µm) were selected per animal, except in cases where tissue integrity was compromised, in which case only two ROIs were used (which occurred in a few sections). The same representative ROIs from each animal and label were used for all analyses.

### 2.8 Intensity analysis

ImageJ software (Fiji) was used for signal intensity analysis. High-resolution images were converted to grayscale and inverted such that strongly positive pixels were displayed in white. The average raw intensity for each ROI was measured using the intensity measure tool in Fiji. All ROIs from a given region were averaged to determine the overall signal intensity of that region (ipsilesional or contralateral ACA, MCA, or striatum). To overcome batch differences in labeling, the ipsilesional values were normalized to the contralateral values within each section to obtain a ratio reflecting the change in the ipsilesional hemisphere signal compared to the expected “normal” signal in the contralateral hemisphere. A ratio of 1.0 indicates no change between hemispheres.

### 2.8 Cell counting

A cloud-based deep learning segmentation platform Biodock (AI Software Platform. Biodock 2024. Available from www.biodock.ai) was used to count the number of cells positive for each marker (GFAP, vimentin, and Iba1). We trained this artificial neural network (ANN) on regions that were not selected for final analysis but were analogous and covered the range of label intensity and cell morphology. An iterative approach was used to accelerate the training process. Initially, a relatively small number of cells were traced by a human observer, these inputs were then used to train a preliminary model, and the predictions of the preliminary model were then further refined by a human expert, which then fed into the final training sets.

We trained three separate models for GFAP, vimentin, and Iba1-labeled cells. Biodock was trained on 1717 (GFAP; across 14 iterations) or 1603 (vimentin; across 11 iterations) labeled astrocytes, or 3258 labeled microglia (Iba1; across 8 iterations). To avoid including small spurious false positives, the output cell counts were further refined based on an area cutoff, which was determined by the typical segmentation area per cell reported by a human expert observer. Cells below the following area were excluded: 39 sq microns (GFAP and Iba1 stains) or 78 sq microns (vimentin). We validated each model by comparing Biodock-generated cell counts to manual counts conducted by three independent blind experts (co-authors TLS, BDM, and AM) on a randomly selected and de-identified subset of test images (Supplemental Figure 1). We concluded training when ≥90% of the difference between the ANN and human experts fell within the 95% confidence interval (SF1, 1.96 ± SD, dotted lines). We reported the density of positive cells as the count for each ROI normalized to the area (cells/mm^2^), averaged for each animal.

### 2.9 Statistical analysis

Infarct sizes were analyzed with a two-sample unpaired t-test for each region (cortex, striatum, and hemisphere) followed by post-hoc Bonferroni-Dunn correction for multiple comparisons.

Intensity ratios were analyzed using a one-sample t-test against the null hypothesis of 1.0 (indicating no difference between hemispheres) for each region. The cell count data were analyzed with a paired (between hemispheres within each section) or unpaired (between groups) t-test. ANN method bias (Supplemental Figure 1) was evaluated using a Bland-Altman plot followed by one-sample t-test against the null hypothesis of calculated bias being 0.0. Significance was defined as p-values below 0.05 unless otherwise indicated. All plots are displayed as mean ± SEM. *p<0.05, **p<0.01, ***p<0.001, ****p<0.0001.

## 3. RESULTS

### 3.1 Optimizing a mild stroke model for chronic studies

We tested MCA occlusion (Figure 1A) durations shorter than 60 min and found that a 30 min MCAo results in a subcortical infarct confined mainly to the striatum while sparing the cortex (Figure 1B,C). The striatal infarct induced by 30 min MCAo (57.4 ± 2.6%) was similar to 60 min MCAo (59.0 ± 7.2%; p=0.99), but the cortical infarct was much smaller (30 min: 6.8 ± 7% vs. 60 min: 34 ± 8.4%; p=0.009), resulting in an overall smaller hemispheric infarction (30 min: 11.18 ± 2% vs. 60 min: 28 ± 5.8%; p=0.0267). We refer to this 30-min MCAo as the mild sub-cortical infarct (MSCI) model.

### 3.2 Reactive astrogliosis spreads throughout the MCA territory in the first week after MSCI

We next examined the extent and progression of reactive astrogliosis induced by MSCI during the first week by immunolabeling for GFAP and vimentin. GFAP is an intermediate filament protein that is expressed largely in astrocytes in the brain and is broadly used as an indicator of reactive astrocytes due to its increased expression following injury. In contrast, vimentin is minimally expressed in healthy mature astrocytes but is upregulated in reactive astrocytes (Escartin et al. 2021). We analyzed GFAP and vimentin in three regions: the ACA cortex (which experiences no blood flow reduction during MSCI), the MCA cortex (which experiences blood flow reduction but no infarction during MSCI), and the striatum (which experiences both blood flow reduction and infarction during MSCI) at 1, 3, and 7 days after MSCI (Figure 1D).

To assess GFAP labeling intensity, we calculated the ratio of ipsilesional to contralateral signal, where a ratio of 1.0 indicates no change. GFAP intensity increased in the MCA cortex and striatum after MSCI but not in the ACA cortex (Figure 2D; Supplementary Table 1). This increase was detectable within 1 day after MSCI and became more pronounced by 7 days. The number of GFAP-positive cells also increased in the ipsilesional MCA cortex and striatum starting 3 days after MSCI and persisted through 7 days (Figure 2F; Supplementary Table 1).

**Figure 2.**
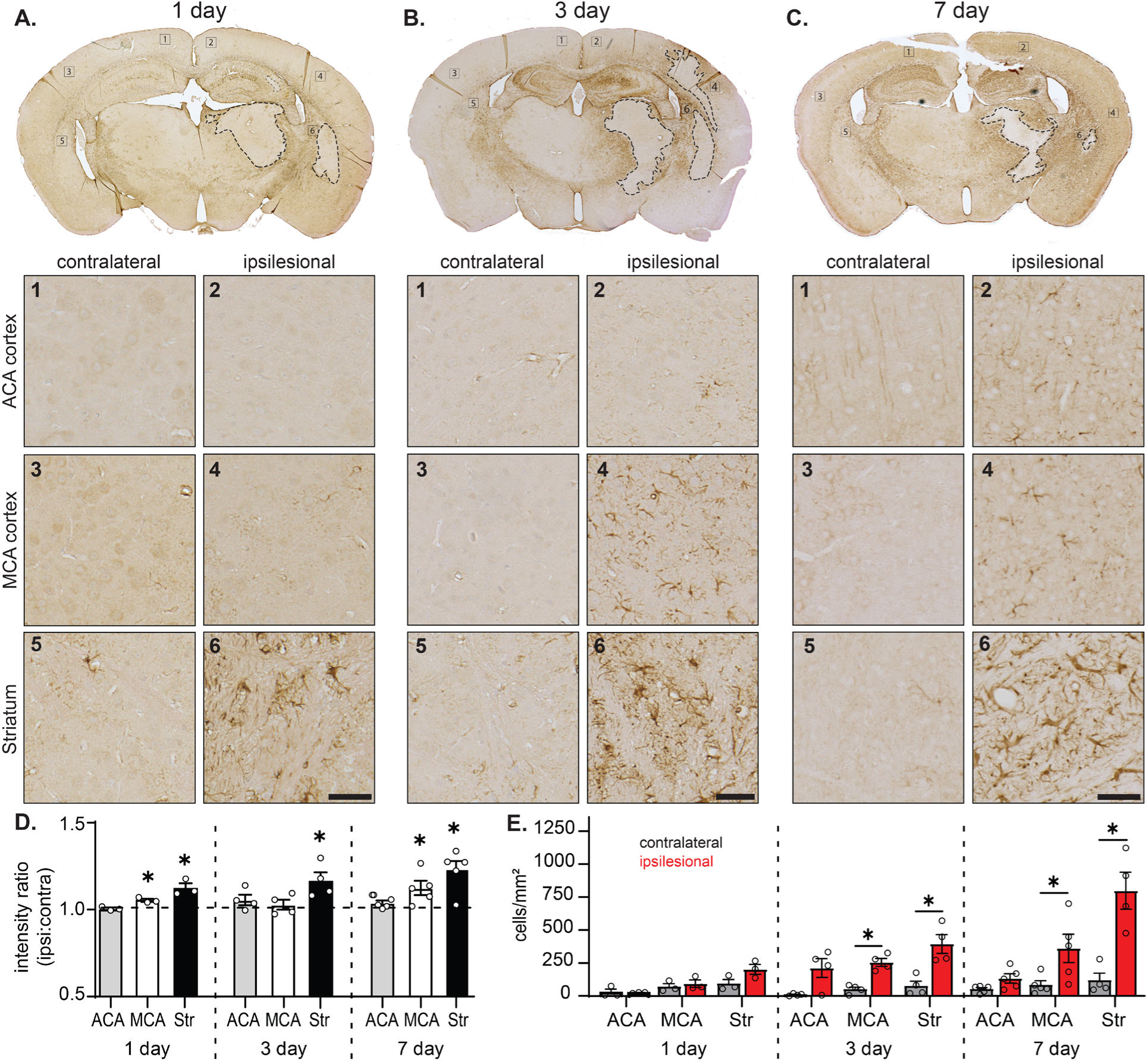
GFAP-labeled astrocytes increase and spread throughout the MCA territory in the sub-acute phase after MSCI. A–C. Top, example coronal sections labeled with GFAP at 1 day (**A**), 3 days (**B**), or 7 days (**C**) after MSCI. Infarct border depicted by dotted line. Bottom, each panel shows high-resolution images of the regions identified by the corresponding number in the top panel coronal section, depicting contralateral and ipsilesional ACA territory, MCA territory, and the non-infarcted border region of the striatum at each timepoint. Scale bars = 50 µm. **D**. Ratio of ipsilesional to contralateral GFAP intensity in the ACA (gray bars), MCA (white bars), and non-infarcted striatum (black bars) at each timepoint after MSCI. **E**. Number of GFAP-positive astrocytes in the contralateral (gray bars) vs. ipsilesional (red bars) hemispheres in the ACA, MCA, and striatum at each timepoint after MSCI. Data shown as mean ± SEM. *p<0.05.

The ipsilesional ACA cortex showed a nonsignificant trend toward an increase in GFAP-positive cells at 3 days after MSCI, but levels returned to baseline by 7 after MSCI.

Vimentin followed a similar pattern. The vimentin labeling intensity ratio (ipsilesional:contralateral) increased in the MCA cortex and striatum b 1 day after MSCI and remained elevated through 7 days, with some variability evident at the 3-day timepoint (Figure 3D; Supplementary Table 1). At day 1, the number of vimentin-positive cells in the ipsilesional hemisphere remained low across all regions, similar to the contralateral hemisphere, but increased in both the MCA cortex and striatum at 3 and 7 days after MSCI (Figure 3F; Supplementary Table 1). No changes were observed in vimentin levels in the ACA region of the cortex.

**Figure 3.**
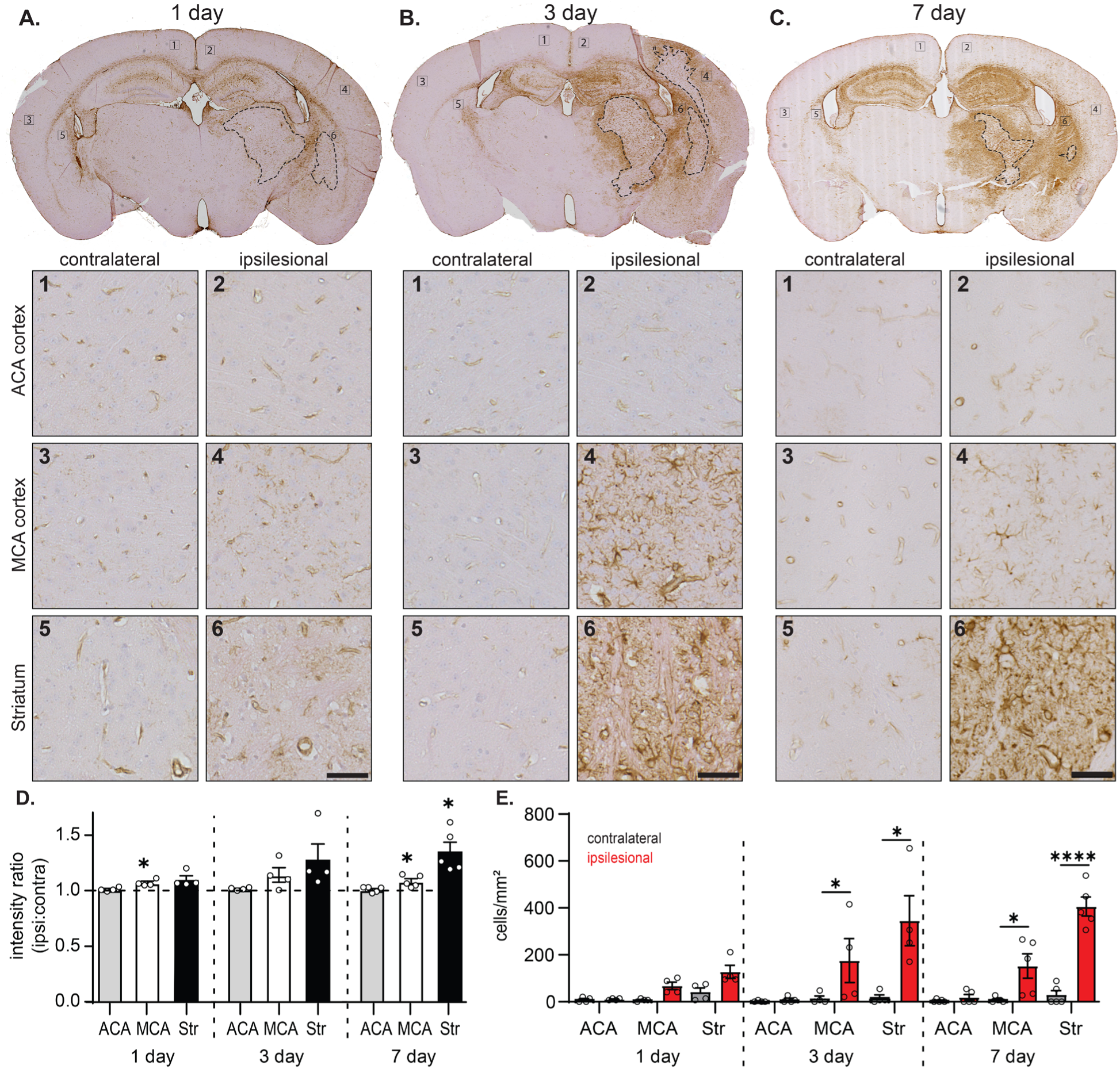
Vimentin-labeled reactive astrocytes increase and spread during the sub-acute period after MSCI. A–C. Top, example coronal sections labeled with vimentin at 1 day (**A**), 3 days (**B**), or 7 days (**C**) after MSCI. Infarct border depicted by dotted line. Bottom, each panel shows high-resolution images of the regions identified by the corresponding number in the top panel coronal section, depicting contralateral and ipsilesional ACA, MCA, and non-infarcted border striatum regions at each timepoint. Scale bars = 50 µm. **D.** Ratio of ipsilesional to contralateral vimentin intensity for each region and timepoint after MSCI. **E.** Number of vimentin-positive astrocytes in the contralateral (gray bars) vs. ipsilesional (red bars) hemispheres for each region and timepoint after MSCI. Data shown as mean ± SEM. *p<0.05, ****p<0.0001.

Together, these data suggest that the reactive astrogliosis response begins within the first day after a small subcortical stroke and progressively increases throughout the sub-acute phase (first week).

### 3.3 Microglia reactivity is restricted near the infarcted region after MSCI

Microglia play a crucial role in injury response and recovery, including after ischemic stroke (Paolicelli et al. 2022; Gao et al. 2023). Microglia migrate to and proliferate at the site of injury; therefore, we considered four regions for analysis: the ACA cortex, the MCA cortex, the striatum border, and the infarcted striatum. The infarct border was identified using H&E staining and confirmed by the absence of GFAP labeling in adjacent sections (dotted lines, Figure 4A–C).

**Figure 4.**
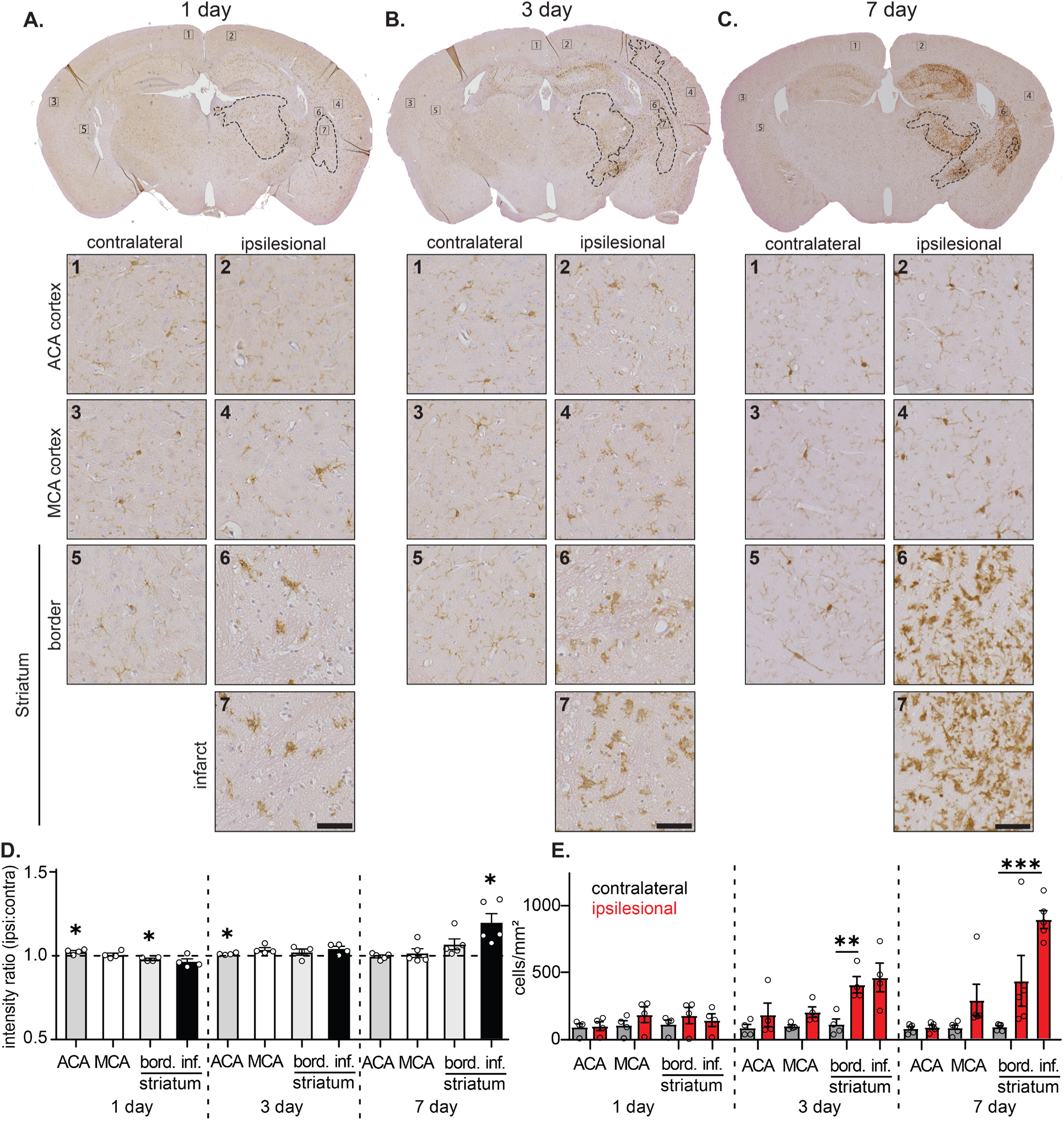
Iba1-labeled microglia increase after MSCI in infarct-adjacent regions. A–C. Top, example coronal sections labeled with Iba1 at 1 day (**A**), 3 days (**B**), or 7 days (**C**) after MSCI. Infarct border depicted by dotted line. Bottom, each panel shows high-resolution images of the regions identified by the corresponding number in the top panel coronal section, depicting contralateral and ipsilesional ACA, MCA, and striatum (both infarct and border regions shown for the ipsilesional hemisphere). Scale bars = 50 µm. **D.** Ratio of the ipsilesional to contralateral Iba1 intensity for each region and timepoint after MSCI. **E.** Number of Iba1-positive astrocytes in the contralateral (gray bars) vs. ipsilesional (red bars) hemispheres for each region and timepoint after MSCI. Data shown as mean ± SEM for all bars. *p<0.05, **p<0.01, ***p<0.001.

The Iba1 intensity ratio (ipsilesional:contralateral) did not increase within the striatal infarct until 7 days after MSCI (Figure 4D; Supplementary Table 1). Although the ACA cortex showed a significant change at 1 day, the change was minimal and attributed to low variability affecting statistical calculations. Accordingly, the number of Iba1-positive microglia remained unchanged in the ACA and MCA cortices but increased within the striatum in the infarct and its surrounding border region, with the highest numbers observed in the infarct at 7 days post-MSCI (Figure 4E; Supplementary Table 1). These data suggest that the microglial reactivity response is largely localized to the infarcted region.

### 3.4 Microglia limit infarct size but are not necessary for reactive astrogliosis after stroke

The delayed and spatially restricted microglia reactivity we observed, in contrast to the widespread reactive astrogliosis, indicates that post-ischemic astrogliosis may not be regulated by microglial signals, as has been shown in other pathologies (Liddelow et al. 2017; Shinozaki et al. 2017). To investigate the relationship between reactive microglia and astrogliosis after stroke, we depleted microglia in our MSCI model using Pexidartinib (PLX3397), a colony-stimulating factor 1 receptor (CSF1R) antagonist. CSF1 regulates the survival, proliferation, and differentiation of monocyte and macrophage lineage cells (Fujiwara et al. 2021; Hagan et al. 2020), and CSF1R inhibition depletes microglia (Elmore et al. 2014). To examine the microglial contribution to astrogliosis after MSCI, we fed mice with chow containing PLX3397 (290 mg/kg) continuously for 3 weeks (Figure 5A).

**Figure 5.**
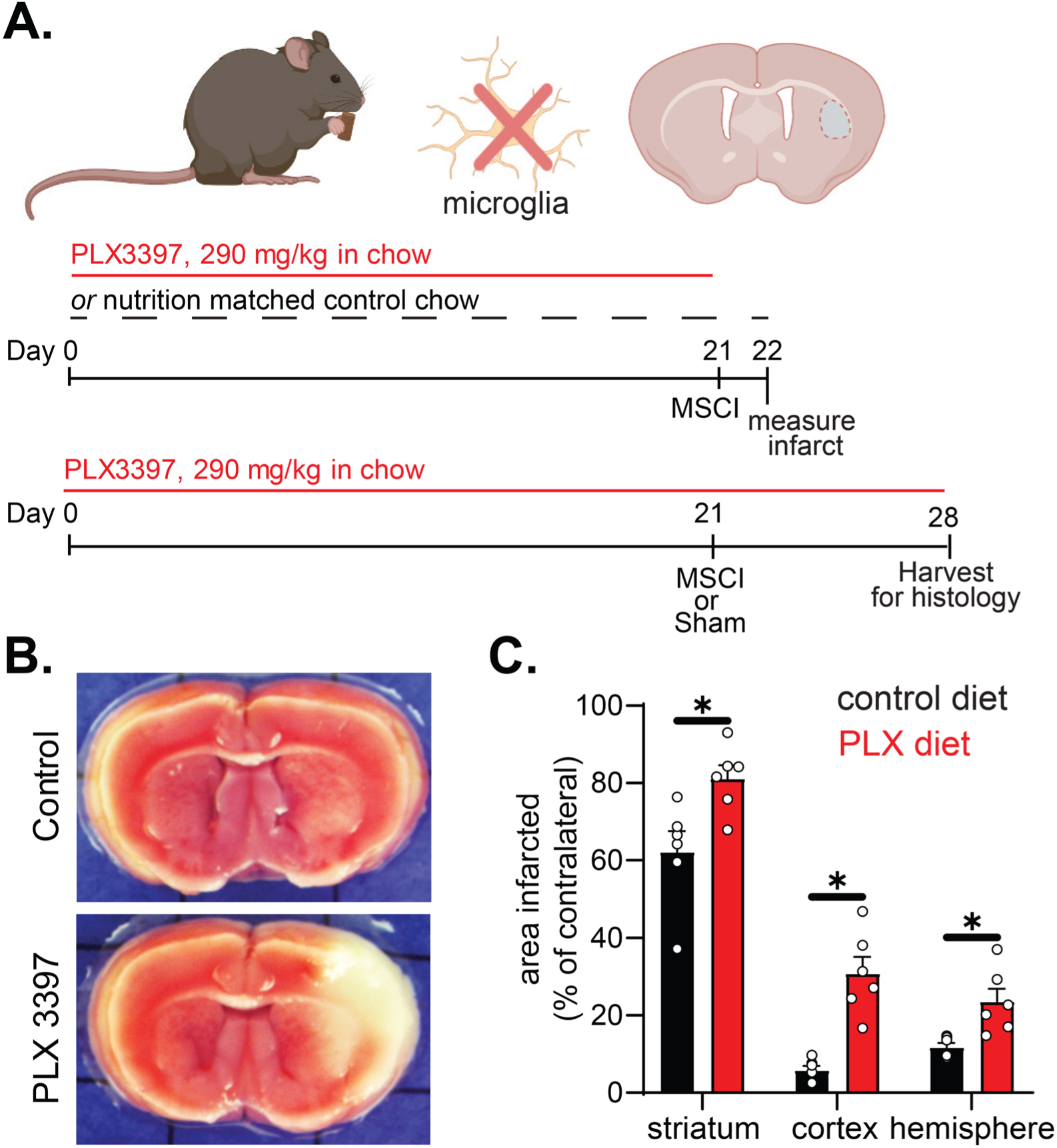
Microglia depletion increases infarct size after MSCI. **A.** Experimental timeline for microglia depletion. Mice were fed chow containing 290 mg/kg PLX3397 or nutrition-matched control chow for 3 weeks. One cohort of mice (n = 6 control diet, 6 PLX diet) received MSCI on day 21 and were sacrificed 24 h later to assess infarct size. A separate cohort (n = 4 sham surgery, n = 6 MSCI) received MSCI or sham surgery and was survived for another 7 days under the same diet before tissue collection for histology. **B.** Example brain sections stained with TTC (infarct is seen as absence of the stain) from animals fed control (top) or PLX3397 (bottom) diet. **C.** PLX3397-treated animals exhibited a larger infarct in the striatum and cortex, constituting a larger region of the hemisphere. Data shown as mean ± SEM for all bars. *p<0.05. Schematic in A created with BioRender.com.

We first observed that infarct size following 30-minute MCAo was significantly larger in mice that were fed PLX3397-impregnated chow, with infarcts extending into the cortex compared to those fed the control diet (Figure 5B,C). PLX3397 treatment drastically depleted microglia (Figure 6C) and eliminated hemispheric differences in Iba1 labeling intensity after MSCI across all regions at the 7-day timepoint (Figure 6D, right panel; Supplementary Table 2). A small increase in Iba1-positive cell count was observed within the infarcted striatum (Figure 6E, right panel; Supplementary Table 2), but these cells were sparse compared to the extensive Iba1 labeling seen in mice with intact microglia at 7 days post-MSCI (Figure 4). Next, we examined reactive astrogliosis at 7 days after MSCI, when astrogliosis was most pronounced in control mice (Figures 2 and 3). We observed an increase in both GFAP and vimentin labeling intensity, as well as a higher number of positive cells, in the ipsilesional MCA and striatum compared to analogous regions in the contralateral hemisphere and animals fed the control diet (Figure 6D–E, left and middle panels; Supplementary Table 2). These data show that microglia are important in limiting the infarct size but that stroke-induced reactive astrogliosis remains largely intact even when microglia are severely depleted.

**Figure 6.**
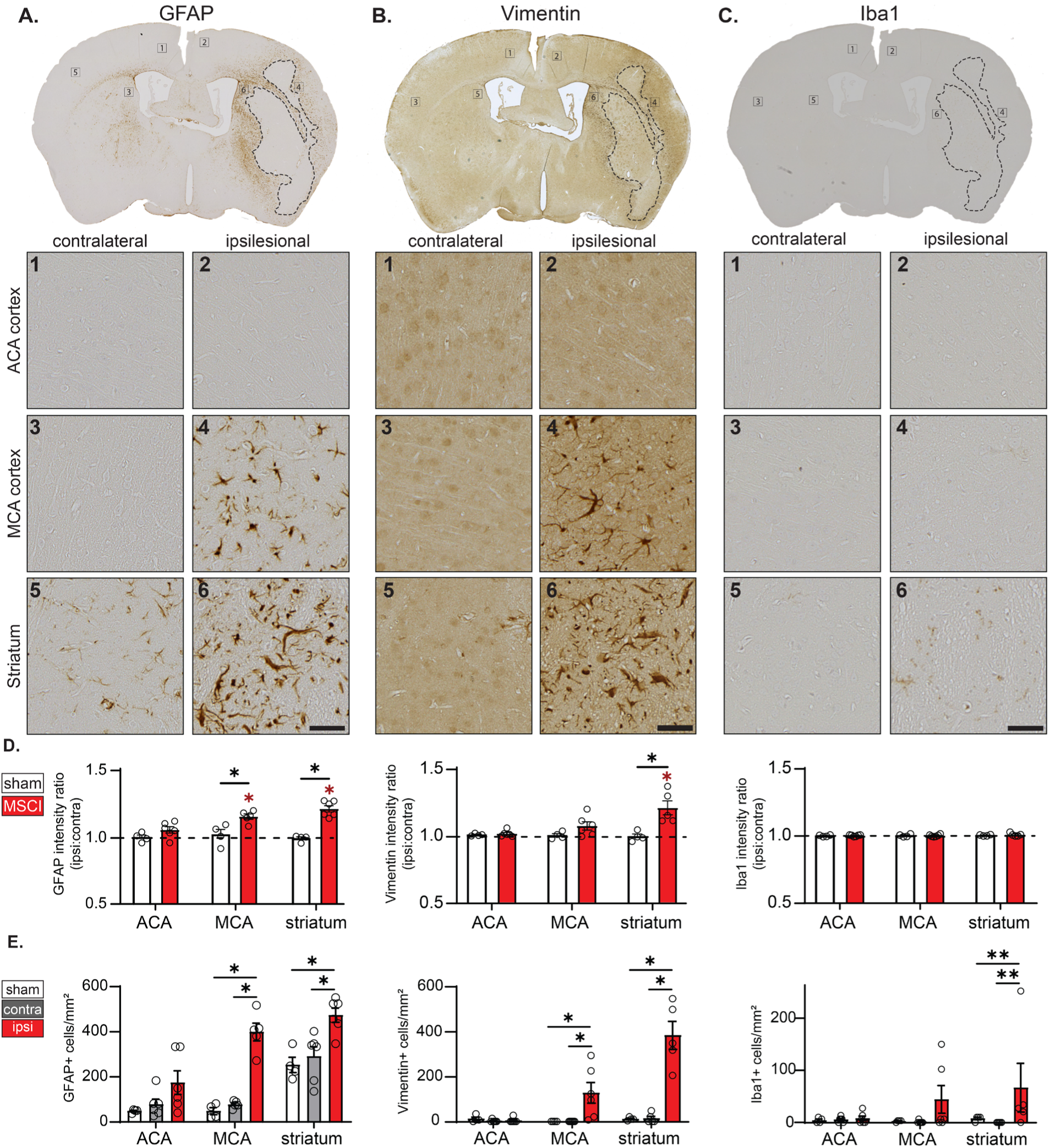
Microglia depletion does not abolish astrogliosis after MSCI. A–C. Top, example coronal sections labeled with GFAP (**A**), vimentin (**B**), or Iba1 (**C**) 7 days after MSCI. Infarct border depicted by dotted line. Bottom, each panel shows high-resolution images of the regions identified by the corresponding number in the top panel coronal section. Scale bars = 50 µm. **D.** Ipsilesional to contralateral intensity ratio of GFAP (left), vimentin (middle), or Iba1 (right) for ACA, MCA, and striatum in sham (white bars) and MSCI (red bars) animals. **E.** Number of cells positive for GFAP (left), vimentin (middle), or Iba1 (right) in sham (white bars) animals, and in the contralateral (gray bars), and ipsilesional (red bars) hemispheres of MSCI animals. Data shown as mean ± SEM for all bars. *p<0.05, **p<0.01.

### 3.5 MSCI-induced reactive astrogliosis and microglial response remain intact in aged animals

Age is a key factor in stroke recovery, with older patients generally experiencing worse outcomes after an ischemic stroke (Benjamin et al. 2018). We examined whether age influences the progression of reactive astrogliosis and microglia responses by inducing MSCI in ∼20-month-old mice. At baseline, we observed a higher number of GFAP-positive cells in both the ACA and MCA cortices of the contralateral hemisphere of aged MSCI mice, as well as in those that underwent a sham procedure, compared to young mice (Supplementary Table 3). At 7 days after MSCI, the GFAP intensity ratio (ipsilesional:contralesional) increased in the ACA and MCA, with the striatum showing a similar trend (Figure 7C). Additionally, GFAP-positive cell counts increased, particularly in the striatum (Figure 7D; Supplementary Table 3). We observed a similar increase in vimentin signal, with intensity increasing in the striatum and cell counts increasing in both the MCA cortex and the striatum (Figure 8C–D). Iba1 labeling also appeared stronger in aging mice at baseline (Supplementary Table 1). At 7 days post-MSCI, Iba1 labeling intensity and the number of Iba1-positive cells increased within the striatal infarct and infarct border (Figure 9C–D). Overall, these data suggest that, despite a higher baseline level of GFAP and Iba1, reactive astrogliosis and microglia response to stroke remain intact in older animals.

**Figure 7.**
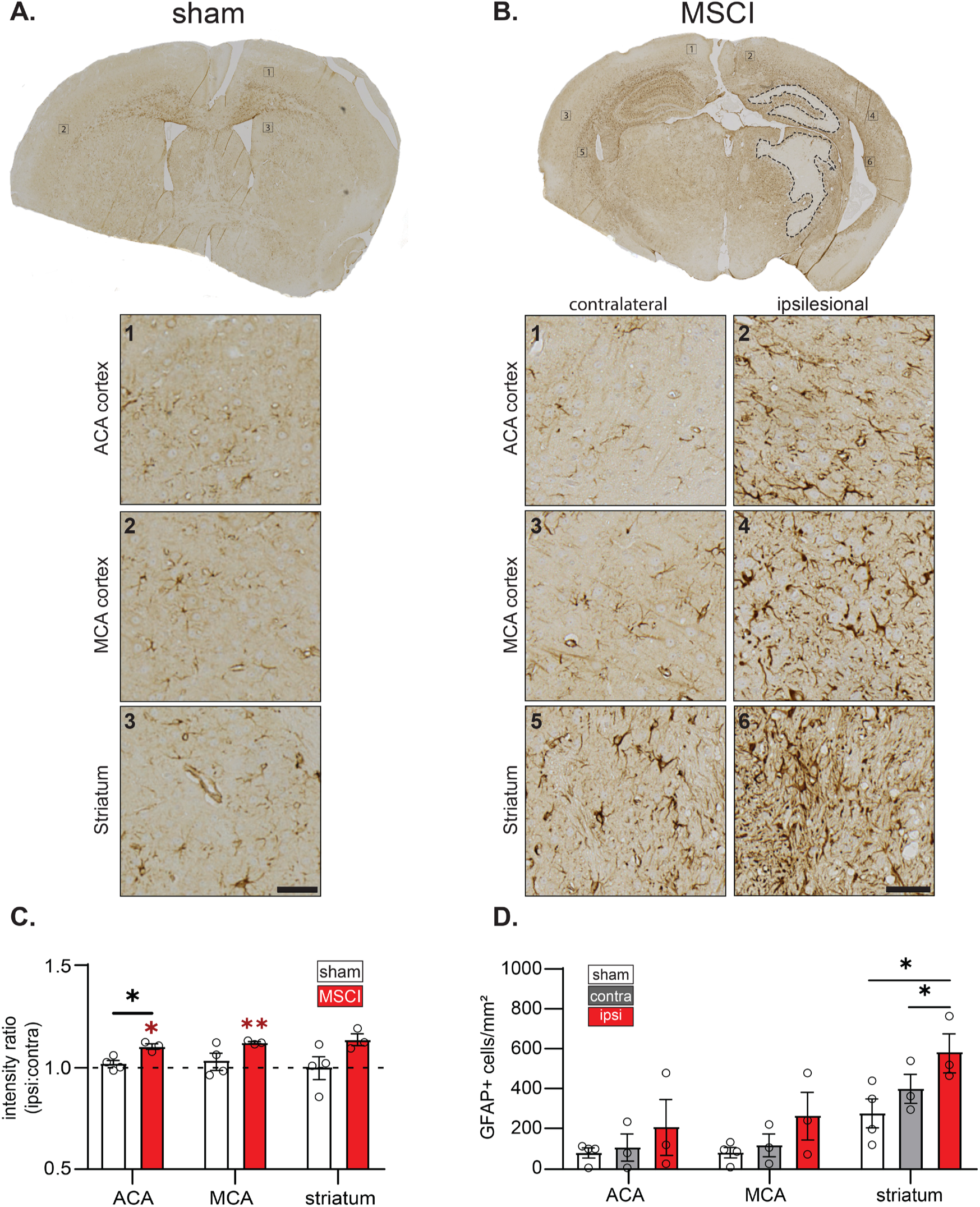
GFAP-labeled astrocytes increase and spread throughout the MCA territory after MSCI in aging animals. A–B. Top, example coronal sections labeled with GFAP 7 days after a sham (**A**) or MSCI (**B**) surgery in 20-month-old animals. Bottom panels show high-resolution images of the regions identified by the corresponding number in the top panel coronal section. Infarct border depicted by dotted line (**B,** MSCI). Scale bars = 50 µm. **C.** Ipsilesional to contralateral intensity ratio of GFAP in MSCI (red bars) or sham (white bars) groups for indicated regions. **D.** Number0 of cells positive for GFAP in the sham animals (white bars), and contralateral (gray bars) and ipsilesional (red bars) hemispheres of MSCI animals by region. Data shown as mean ± SEM for all bars. *p<0.05.

**Figure 8.**
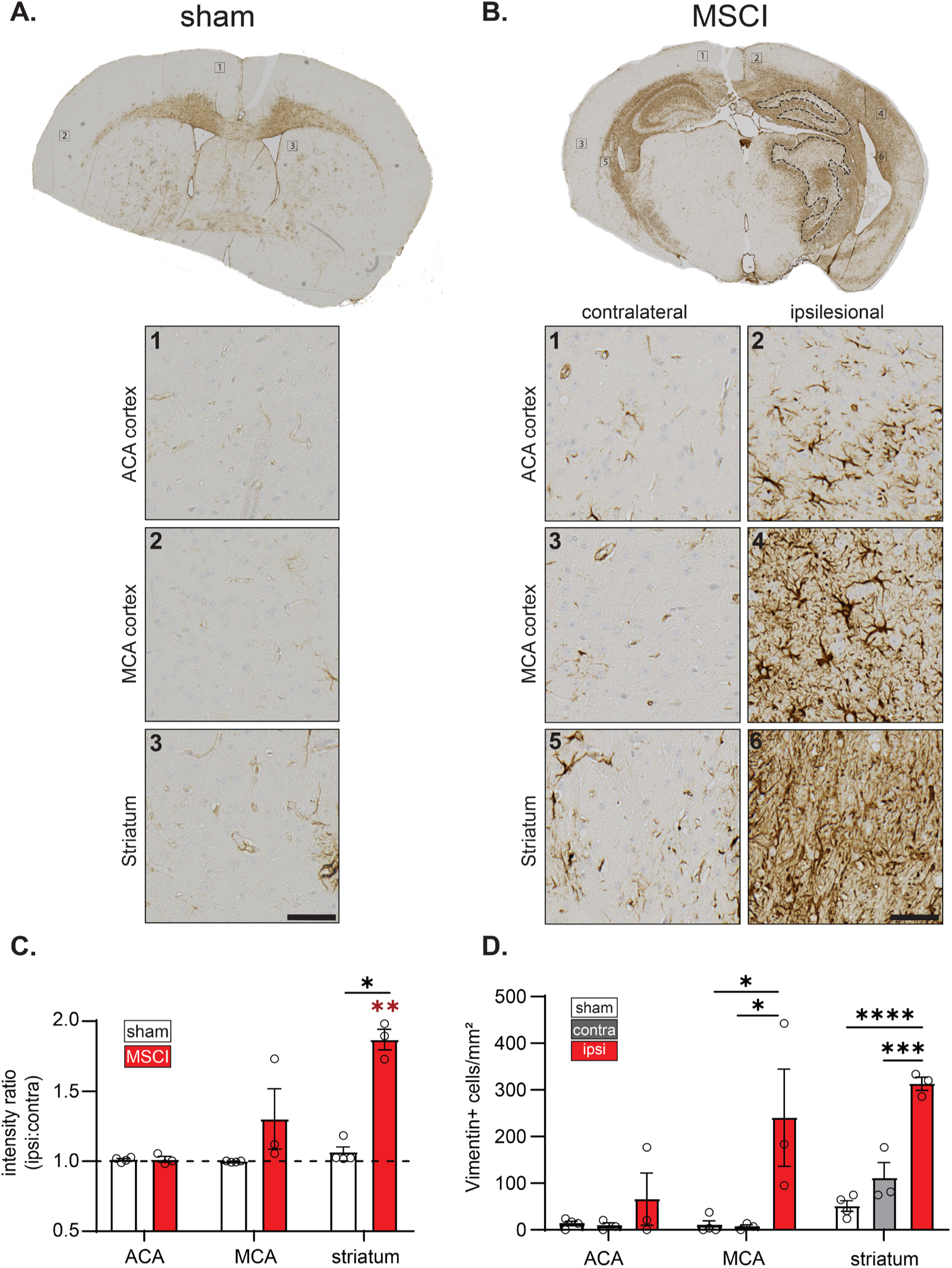
Vimentin-labeled reactive astrocytes also increase and spread after MSCI in aging animals. **A–B.** Top, example coronal sections labeled with vimentin 7 days after a sham (**A**) or MSCI (**B**) surgery in 20-month-old animals. Bottom panels show high-resolution images of the regions identified by the corresponding number in the top panel coronal section. Infarct border depicted by dotted line (**B,** MSCI). Scale bars = 50 µm. **C.** Ipsilesional to contralateral intensity ratio of vimentin in MSCI (red bars) or sham (white bars) animals for each indicated region. **D.** Number of cells positive for vimentin in sham animals (white bars), or contralateral (gray bars) and ipsilesional (red bars) hemispheres of MSCI animals by region. Data shown as mean ± SEM for all bars. *p<0.05, **p<0.01, ***p<0.001, ****p<0.0001.

**Figure 9.**
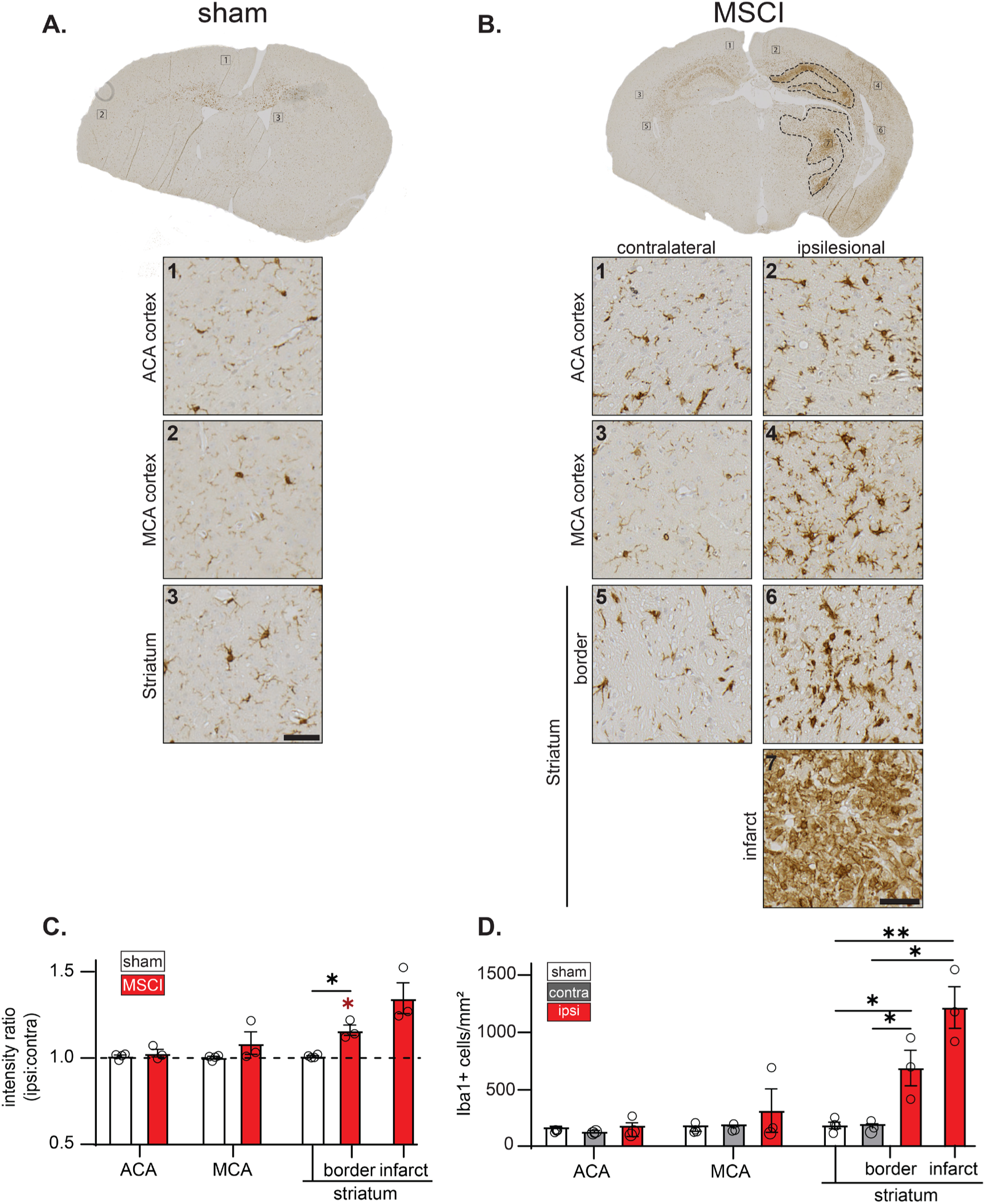
Iba1-labeled microglia increase after MSCI in infarct-adjacent areas in aging animals. **A–B.** Top, example coronal sections labeled with Iba1 7 days after a sham (**A**) or MSCI (**B**) surgery in 20-month-old animals. Bottom panels show high-resolution images of the regions identified by the corresponding number in the top panel coronal section. Infarct border depicted by dotted line (**B,** MSCI). Scale bars = 50 µm. **C.** Ipsilesional to contralateral intensity ratio of Iba1 for MSCI (red bars) or sham (white bars) animals for each indicated region. **D.** Number of cells positive for Iba1 in sham animals (white bars), or contralateral (gray bars) and ipsilesional (red bars) hemispheres of MSCI animals by region. Data shown as mean ± SEM for all bars. *p<0.05, **p<0.01.

## 4. DISCUSSION

We report an optimized model of MCAo with a reduced occlusion period of 30 minutes. This model provides a clinically relevant small subcortical infarct (Figure 1) for studying the effects of stroke in regions distant from the infarct, including an intact cortex. The MCA cortex is particularly relevant, as it experiences decreased blood flow without infarction, which may represent a model of transient ischemic attack and lead to pathologies associated with stroke-related cognitive decline over time. As mice recover well from this shorter occlusion, this model shows promise for exploring chronic post-stroke outcomes. Additionally, provided there is careful control of age and other variables that influence infarct size, it may be suitable for studying striatal-specific strokes while minimizing extensive extra-striatal infarcts.

Our analysis of gliosis using this small stroke model demonstrates that reactive astrogliosis is more pronounced at earlier timepoints after minor ischemic injuries compared to microglia reactivity (Figures 2–4). More importantly, reactive astrogliosis is more widespread, extending into the MCA region of the cortex, whereas microglia reactivity remains largely restricted to the area of infarct and its surrounding border regions. Additionally, despite the physical proximity of MCA and ACA regions, reactive astrogliosis in the cortex remains largely restricted to the MCA territory during the sub-acute phase, suggesting that reactive astrogliosis may primarily be a response to the ischemic injury itself rather than by diffusible signals released from the injured tissue. Future long-term studies may shed light on whether diffusible signals contribute to the spread of gliosis at more chronic timepoints. The widespread nature of reactive astrogliosis, especially at later sub-acute timepoints, raises many new questions. How are astrocyte functions encompassing neuronal circuit modulation, metabolic and ionic homeostasis, and cerebrovascular regulation altered in these distal regions after a small localized ischemic injury, and how do they evolve over time? These are important avenues for future research.

Our observations may not encompass all reactive astrocyte populations, including any population not immunoreactive for vimentin or GFAP (Escartin et al. 2021). Reactive astrogliosis is a complex process involving morphological, transcriptional, translational, and functional changes in astrocytes (Escartin et al. 2021). Thus, relying on a single marker provides only a limited understanding of astrogliosis. Nevertheless, immunolabeling for specific markers represents a first step in identifying distant brain regions potentially affected by subcortical stroke or other injuries for further functional investigations. Although GFAP is widely used as a marker for reactive astrocytes, its baseline expression is naturally higher in some regions (e.g., white matter, hippocampus) compared to others (e.g., cortex) (Kimelberg 2004) and can increased due to certain physiological activities (Escartin et al. 2021). We suggest that vimentin, which is low in mature grey matter astrocytes but highly induced following injury (Figure 3), may serve as a more reliable reporter than GFAP for assessing astrogliosis during acute and sub-acute phases.

In our study, Iba1 labeling did not immediately increase within the infarcted region 1 day after stroke but was robustly observable at later timepoints. This was unexpected, as previous studies have reported rapid recruitment and proliferation of microglia at the site of infarct or ischemic injury (Huang et al. 2023; Hu et al. 2012). Several factors could explain this discrepancy, including the smaller stroke size in our model or the possibility that microglia responding to the ischemic injury do not immediately increase Iba1 expression. Indeed, Iba1 labeling showed less ramified, more amoeboid cells in the striatal infarct and border regions even 1 day after stroke (Figure 4A). Given that Iba1 is not the only marker of reactive microglia, other populations might be overlooked in this study. Further, blood-brain barrier impairment in the infarct leads to blood monocytes infiltrating the brain, where they become macrophages that adopt microglial identity and properties (Han, Liu, and Gao 2020). The large increase in Iba1 positive cell population likely reflects both locally migrating microglia and these infiltrating macrophages, which are also not discerned in our study.

The striking increase in infarct size in mice depleted of microglia (Figure 5), similar to previous findings (Huang et al. 2023), indicates that microglia are crucial for maintaining the infarct border and limiting injury. Microglia are thought to be necessary for the induction of a border (scar) forming reactive astrocyte subtypes, which protect against neurotoxic inflammation (Sofroniew 2015). Although we observed largely intact reactive astrogliosis after microglia depletion (Figure 6), their ability to form the injury border may still be impaired. The absence of other key microglial functions, such as debris clearance, reactive oxygen species sequestration, or inflammatory signaling regulation (Xiong, Liu, and Yang 2016; Wang, Leak, and Cao 2022), might also contribute to the increased infarct size. Importantly, our observation that reactive astrogliosis persists in microglia-depleted mice suggests that its extent is not solely dependent on signals from microglia in ischemic stroke, unlike in post-infection contexts (Liddelow et al. 2017). Instead, astrogliosis in this model may be influenced directly by factors such as reduced blood flow during stroke, hypoxia, metabolic changes, oxidative stress, or signals released from injured tissue. Single-cell analyses have identified multiple subclusters of reactive astrocytes in the days and weeks following a stroke (Guo et al. 2021; Zhang et al. 2022). It is possible that only certain subsets of reactive astrocytes are dependent on signals from microglia, while others, as observed here, are not.

Reactive astrogliosis and reactive microglia may be “primed” for increased reactivity based on many environmental factors, including age. Indeed, we observed an increase in GFAP and Iba1 in aged mice even at baseline (although direct statistical comparisons were not possible as the tissues were processed in separate batches). This finding generally aligns with previous characterizations of astrocytes (Clarke et al. 2018; Suda et al. 2021) and microglia (Suda et al. 2021) in aging. Despite these baseline differences, the MSCI-induced increase in both reactive astrocytes and microglia, as well as their spatiotemporal patterns, remained similar between aged mice and middle-aged adult mice (Figures 7–9). Age could also impact the size of the infarct, which in turn could affect the extent of reactive astrogliosis or microglia reactivity. Although infarct size in aged mice was not quantified in this study, we observed that one of the aged mice exhibited an infarct that spread even to the hippocampus (example shown in panel B of Figures. 7–9), an area that is normally supplied by the posterior cerebral artery rather than the MCA. Whether this is due to aging itself or whether this particular mouse had a misformed artery structure in which the MCA supplied the hippocampus is unknown. A related limitation of our study is that all experiments were conducted in C57/B6 mice, in which the circle of Willis can be variable (Doyle et al. 2012). This variability may have contributed to the larger infarct in this particular mouse. The effect of age on the impact of stroke presents an intriguing avenue for future research, as understanding the relationship between age, infarct size, and reactive glial responses could yield significant insights into stroke pathology and recovery, as well as the underlying causes of stroke-induced dementia.

## Conclusion

Here we describe a MSCI model suitable for chronic studies on the effects of ischemic stroke. We identified that, in the sub-acute phase (up to 7 days after MSCI), reactive astrogliosis precedes reactive microglia, and this response is restricted primarily to the arterial territory that experienced ischemia, with or without infarction. We found that stroke-induced reactive astrogliosis is, to a great extent, not dependent on microglia. Finally, we show that although aging is associated with some increase in astrocyte and microglia markers, the response of both cell types to stroke is conserved, at least during the sub-acute period. Our findings raise important questions for future research: what signals induce astrocyte and microglia reactivity after ischemic stroke, and how do they diverge? How long do reactive astrogliosis and microglia persist, and could there be permanent astrocyte or microglia state changes after injury? And perhaps most importantly, what are the functional consequences of widespread reactive gliosis during the sub-acute and chronic phases, and how do they contribute to neurodegeneration, such as in post-stroke dementia?

## Author contributions

Experimental design by AM, TLS, and HLM. Data collection by TLS, BDM, SJS, HLM, LKM, RDLT, AH, RW and AM. MCAo surgery by WZ. Data analysis by TLS. Manuscript writing by TLS and AM and editing by all co-authors.

## Data Availability Statement

The data presented in this study are available upon reasonable request.

## Ethics Statement

The animal study was reviewed and approved by the Oregon Health & Science University Institutional Animal Care and Use Committee.

## Acknowledgments

We acknowledge the OHSU Advanced Light Microscopy Core (RRID:SCR_009961) for providing expert technical assistance and Dr. Laura Villasana for providing mice for the aging experiments. This work was funded by a Lacroute fellowship (TLS), JTMF Foundation award (AM) and several NIH grants: NIA NRSA T32AG055378 (TLS), NHLBI NRSA T32HL094294 (HM), NINDS R01NS110690 and R01NS134592 (AM), and NIA P30AG066518 (AM, RW; Program directors: Silbert, Lisa & Lim, Miranda).

## Supplementary figure legend

**Supplemental Figure 1. Trained artificial neural networks reliably detect astrocytes and microglia. A-C.** Representative examples of original images and artificial neural net (ANN)-identified cell outlines for the same images. Two example morphological profiles of astrocytes with GFAP labeling (A) or vimentin labeling (B) and microglia with Iba1 labeling (C) are shown to display the range of morphology the ANN can identify. The vimentin images show how the ANN specifically detected astrocytes but not vascular structures, which are also positive for vimentin (B). **D-F.** Bland-Altman plots showing the difference in cell counts reported by the ANN vs. human experts (averaged counts from three humans: co-authors TLS, BDM, and AM) on the y-axis compared to the average of the ANN and human count on the x-axis. A subset of analysis regions (10 each for GFAP and Vimentin, 12 for Iba1) were used to validate each label. Red line depicts the calculated bias. A one-sample t-test against 0.0 (indicating no bias) shows that no significant bias was detected for any label. We concluded training when ≥90% of the difference between the ANN and human experts fell within the 95% confidence interval (1.96±SD, dotted lines on Bland-Altman plots).

## Supplementary table legends

**Supplementary Table 1. Summary data values for GFAP, vimentin, and Iba1 labels at 1, 3, and 7-days post-MSCI.** The mean intensity ratio and cell counts for each immunolabel are listed by region and hemisphere, as appropriate, for 6-month-old mice exposed to MSCI. N corresponds to number of independent animals used. Data shown as mean ± SEM. Data for GFAP corresponds to Figure 2, vimentin to Figure 3, and Iba1 to Figure 4.

**Supplementary Table 2. Summary data values for GFAP, vimentin, and Iba1 immunolabels after PLX3397-driven microglia depletion.** The mean intensity ratio and cell counts for each immunolabel are listed by region and hemisphere (cell count values only, MSCI) for MSCI or sham 6-month-old mice. N corresponds to number of independent animals used per group. Data shown as mean ± SEM. Data corresponds to Figure 6.

**Supplementary Table 3. Summary data for aging MSCI experiments.** The mean intensity ratio and cell counts for each immunolabel (GFAP, vimentin, Iba1) are listed by region and hemisphere, as appropriate for MSCI or sham 20-month-old mice. N corresponds to the number of independent animals used per group. Data for GFAP corresponds to Figure 7, vimentin to Figure 8, and Iba1 to Figure 9.

## References

1. Amantea, D., G. Micieli, C. Tassorelli, M. I. Cuartero, I. Ballesteros, M. Certo, M. A. Moro, I. Lizasoain, and G. Bagetta. 2015. ‘Rational modulation of the innate immune system for neuroprotection in ischemic stroke’, Front Neurosci, 9: 147.

2. Anderson, M. A., J. E. Burda, Y. Ren, Y. Ao, T. M. O’Shea, R. Kawaguchi, G. Coppola, B. S. Khakh, T. J. Deming, and M. V. Sofroniew. 2016. ‘Astrocyte scar formation aids central nervous system axon regeneration’, Nature, 532: 195–200.

3. Barreto, G.E., X. Sun, L. Xu, and R.G. Giffard. 2011. ‘Astrocyte proliferation following stroke in the mouse depends on distance from the infarct’, PLoS.One., 6: e27881.

4. Barthels, D., and H. Das. 2020. ‘Current advances in ischemic stroke research and therapies’, Biochim Biophys Acta Mol Basis Dis, 1866: 165260.

5. Benjamin, E. J., S. S. Virani, C. W. Callaway, A. M. Chamberlain, A. R. Chang, S. Cheng, S. E. Chiuve, M. Cushman, F. N. Delling, R. Deo, S. D. de Ferranti, J. F. Ferguson, M. Fornage, C. Gillespie, C. R. Isasi, M. C. Jimenez, L. C. Jordan, S. E. Judd, D. Lackland, J. H. Lichtman, L. Lisabeth, S. Liu, C. T. Longenecker, P. L. Lutsey, J. S. Mackey, D. B. Matchar, K. Matsushita, M. E. Mussolino, K. Nasir, M. O’Flaherty, L. P. Palaniappan, A. Pandey, D. K. Pandey, M. J. Reeves, M. D. Ritchey, C. J. Rodriguez, G. A. Roth, W. D. Rosamond, U. K. A. Sampson, G. M. Satou, S. H. Shah, N. L. Spartano, D. L. Tirschwell, C. W. Tsao, J. H. Voeks, J. Z. Willey, J. T. Wilkins, J. H. Wu, H. M. Alger, S. S. Wong, P. Muntner, Epidemiology American Heart Association Council on, Committee Prevention Statistics, and Subcommittee Stroke Statistics. 2018. ‘Heart Disease and Stroke Statistics-2018 Update: A Report From the American Heart Association’, Circulation, 137: e67–e492.

6. Birenbaum, D., L. W. Bancroft, and G. J. Felsberg. 2011. ‘Imaging in acute stroke’, West J Emerg Med, 12: 67–76.

7. Bisht, K., K. A. Okojie, K. Sharma, D. H. Lentferink, Y. Y. Sun, H. R. Chen, J. O. Uweru, S. Amancherla, Z. Calcuttawala, A. B. Campos-Salazar, B. Corliss, L. Jabbour, J. Benderoth, B. Friestad, W. A. Mills, 3rd, B. E. Isakson, M. E. Tremblay, C. Y. Kuan, and U. B. Eyo. 2021. ‘Capillary-associated microglia regulate vascular structure and function through PANX1-P2RY12 coupling in mice’, Nat Commun, 12: 5289.

8. Burda, J.E., and M.V. Sofroniew. 2014. ‘Reactive gliosis and the multicellular response to CNS damage and disease’, Neuron, 81: 229–48.

9. Carmichael, S.T. 2005. ‘Rodent models of focal stroke: size, mechanism, and purpose’, NeuroRx., 2: 396–409.

10. Clarke, L. E., S. A. Liddelow, C. Chakraborty, A. E. Munch, M. Heiman, and B. A. Barres. 2018. ‘Normal aging induces A1-like astrocyte reactivity’, Proc Natl Acad Sci U S A, 115: E1896–E905.

11. Doyle, K. P., N. Fathali, M. R. Siddiqui, and M. S. Buckwalter. 2012. ‘Distal hypoxic stroke: a new mouse model of stroke with high throughput, low variability and a quantifiable functional deficit’, J Neurosci Methods, 207: 31–40.

12. Elmore, M. R., A. R. Najafi, M. A. Koike, N. N. Dagher, E. E. Spangenberg, R. A. Rice, M. Kitazawa, B. Matusow, H. Nguyen, B. L. West, and K. N. Green. 2014. ‘Colony-stimulating factor 1 receptor signaling is necessary for microglia viability, unmasking a microglia progenitor cell in the adult brain’, Neuron, 82: 380–97.

13. Escartin, C., E. Galea, A. Lakatos, J. P. O’Callaghan, G. C. Petzold, A. Serrano-Pozo, C. Steinhauser, A. Volterra, G. Carmignoto, A. Agarwal, N. J. Allen, A. Araque, L. Barbeito, A. Barzilai, D. E. Bergles, G. Bonvento, A. M. Butt, W. T. Chen, M. Cohen-Salmon, C. Cunningham, B. Deneen, B. De Strooper, B. Diaz-Castro, C. Farina, M. Freeman, V. Gallo, J. E. Goldman, S. A. Goldman, M. Gotz, A. Gutierrez, P. G. Haydon, D. H. Heiland, E. M. Hol, M. G. Holt, M. Iino, K. V. Kastanenka, H. Kettenmann, B. S. Khakh, S. Koizumi, C. J. Lee, S. A. Liddelow, B. A. MacVicar, P. Magistretti, A. Messing, A. Mishra, A. V. Molofsky, K. K. Murai, C. M. Norris, S. Okada, S. H. R. Oliet, J. F. Oliveira, A. Panatier, V. Parpura, M. Pekna, M. Pekny, L. Pellerin, G. Perea, B. G. Perez-Nievas, F. W. Pfrieger, K. E. Poskanzer, F. J. Quintana, R. M. Ransohoff, M. Riquelme-Perez, S. Robel, C. R. Rose, J. D. Rothstein, N. Rouach, D. H. Rowitch, A. Semyanov, S. Sirko, H. Sontheimer, R. A. Swanson, J. Vitorica, I. B. Wanner, L. B. Wood, J. Wu, B. Zheng, E. R. Zimmer, R. Zorec, M. V. Sofroniew, and A. Verkhratsky. 2021. ‘Reactive astrocyte nomenclature, definitions, and future directions’, Nat Neurosci, 24: 312–25.

14. Feske, S. K. 2021. ‘Ischemic Stroke’, Am J Med, 134: 1457–64.

15. Fujiwara, T., M. A. Yakoub, A. Chandler, A. B. Christ, G. Yang, O. Ouerfelli, V. K. Rajasekhar, A. Yoshida, H. Kondo, T. Hata, H. Tazawa, Y. Dogan, M. A. S. Moore, T. Fujiwara, T. Ozaki, E. Purdue, and J. H. Healey. 2021. ‘CSF1/CSF1R Signaling Inhibitor Pexidartinib (PLX3397) Reprograms Tumor-Associated Macrophages and Stimulates T-cell Infiltration in the Sarcoma Microenvironment’, Mol Cancer Ther, 20: 1388–99.

16. Gao, C., J. Jiang, Y. Tan, and S. Chen. 2023. ‘Microglia in neurodegenerative diseases: mechanism and potential therapeutic targets’, Signal Transduct Target Ther, 8: 359.

17. Gong, Z., J. Guo, B. Liu, Y. Guo, C. Cheng, Y. Jiang, N. Liang, M. Hu, T. Song, L. Yang, H. Li, H. Zhang, X. Zong, Q. Che, and N. Shi. 2023. ‘Mechanisms of immune response and cell death in ischemic stroke and their regulation by natural compounds’, Front Immunol, 14: 1287857.

18. Guo, K., J. Luo, D. Feng, L. Wu, X. Wang, L. Xia, K. Tao, X. Wu, W. Cui, Y. He, B. Wang, Z. Zhao, and Z. Zhang. 2021. ‘Single-Cell RNA Sequencing With Combined Use of Bulk RNA Sequencing to Reveal Cell Heterogeneity and Molecular Changes at Acute Stage of Ischemic Stroke in Mouse Cortex Penumbra Area’, Front Cell Dev Biol, 9: 624711.

19. Hachinski, V. 2018. ‘The convergence of stroke and dementia’, Arq Neuropsiquiatr, 76: 849–52.

20. Hagan, N., J. L. Kane, D. Grover, L. Woodworth, C. Madore, J. Saleh, J. Sancho, J. Liu, Y. Li, J. Proto, M. Zelic, A. Mahan, M. Kothe, A. A. Scholte, M. Fitzgerald, B. Gisevius, A. Haghikia, O. Butovsky, and D. Ofengeim. 2020. ‘CSF1R signaling is a regulator of pathogenesis in progressive MS’, Cell Death Dis, 11: 904.

21. Han, D., H. Liu, and Y. Gao. 2020. ‘The role of peripheral monocytes and macrophages in ischemic stroke’, Neurol Sci, 41: 3589–607.

22. Hickman, S., S. Izzy, P. Sen, L. Morsett, and J. El Khoury. 2018. ‘Microglia in neurodegeneration’, Nat Neurosci, 21: 1359–69.

23. Hu, X., P. Li, Y. Guo, H. Wang, R. K. Leak, S. Chen, Y. Gao, and J. Chen. 2012. ‘Microglia/macrophage polarization dynamics reveal novel mechanism of injury expansion after focal cerebral ischemia’, Stroke, 43: 3063–70.

24. Huang, X., M. Guo, Y. Zhang, J. Xie, R. Huang, Z. Zuo, P. E. Saw, and M. Cao. 2023. ‘Microglial IL-1RA ameliorates brain injury after ischemic stroke by inhibiting astrocytic CXCL1-mediated neutrophil recruitment and microvessel occlusion’, Glia, 71: 1607–25.

25. Jia, J., L. Yang, Y. Chen, L. Zheng, Y. Chen, Y. Xu, and M. Zhang. 2021. ‘The Role of Microglial Phagocytosis in Ischemic Stroke’, Front Immunol, 12: 790201.

26. Kimelberg, H. K. 2004. ‘The problem of astrocyte identity’, Neurochem Int, 45: 191–202.

27. Levine, D. A., A. T. Galecki, K. M. Langa, F. W. Unverzagt, M. U. Kabeto, B. Giordani, and V. G. Wadley. 2015. ‘Trajectory of Cognitive Decline After Incident Stroke’, JAMA, 314: 41–51.

28. Li, Z., H. L. McConnell, T. L. Stackhouse, M. M. Pike, W. Zhang, and A. Mishra. 2021. ‘Increased 20-HETE Signaling Suppresses Capillary Neurovascular Coupling After Ischemic Stroke in Regions Beyond the Infarct’, Front Cell Neurosci, 15: 762843.

29. Liddelow, S. A., K. A. Guttenplan, L. E. Clarke, F. C. Bennett, C. J. Bohlen, L. Schirmer, M. L. Bennett, A. E. Munch, W. S. Chung, T. C. Peterson, D. K. Wilton, A. Frouin, B. A. Napier, N. Panicker, M. Kumar, M. S. Buckwalter, D. H. Rowitch, V. L. Dawson, T. M. Dawson, B. Stevens, and B. A. Barres. 2017. ‘Neurotoxic reactive astrocytes are induced by activated microglia’, Nature, 541: 481–87.

30. Longa, E. Z., P. R. Weinstein, S. Carlson, and R. Cummins. 1989. ‘Reversible middle cerebral artery occlusion without craniectomy in rats’, Stroke, 20: 84–91.

31. Mijajlovic, M. D., A. Pavlovic, M. Brainin, W. D. Heiss, T. J. Quinn, H. B. Ihle-Hansen, D. M. Hermann, E. B. Assayag, E. Richard, A. Thiel, E. Kliper, Y. I. Shin, Y. H. Kim, S. Choi, S. Jung, Y. B. Lee, O. Sinanovic, D. A. Levine, I. Schlesinger, G. Mead, V. Milosevic, D. Leys, G. Hagberg, M. H. Ursin, Y. Teuschl, S. Prokopenko, E. Mozheyko, A. Bezdenezhnykh, K. Matz, V. Aleksic, D. Muresanu, A. D. Korczyn, and N. M. Bornstein. 2017. ‘Post-stroke dementia – a comprehensive review’, BMC Med, 15: 11.

32. Miklossy, J. 2003. ’Cerebral hypoperfusion induces cortical watershed microinfarcts which may further aggravate cognitive decline in Alzheimer’s disease’, Neurol Res, 25: 605–10.

33. Mishra, A., G. R. Gordon, B. A. MacVicar, and E. A. Newman. 2024. ‘Astrocyte Regulation of Cerebral Blood Flow in Health and Disease’, Cold Spring Harb Perspect Biol, 16.

34. Paolicelli, R. C., A. Sierra, B. Stevens, M. E. Tremblay, A. Aguzzi, B. Ajami, I. Amit, E. Audinat, I. Bechmann, M. Bennett, F. Bennett, A. Bessis, K. Biber, S. Bilbo, M. Blurton-Jones, E. Boddeke, D. Brites, B. Brone, G. C. Brown, O. Butovsky, M. J. Carson, B. Castellano, M. Colonna, S. A. Cowley, C. Cunningham, D. Davalos, P. L. De Jager, B. de Strooper, A. Denes, B. J. L. Eggen, U. Eyo, E. Galea, S. Garel, F. Ginhoux, C. K. Glass, O. Gokce, D. Gomez-Nicola, B. Gonzalez, S. Gordon, M. B. Graeber, A. D. Greenhalgh, P. Gressens, M. Greter, D. H. Gutmann, C. Haass, M. T. Heneka, F. L. Heppner, S. Hong, D. A. Hume, S. Jung, H. Kettenmann, J. Kipnis, R. Koyama, G. Lemke, M. Lynch, A. Majewska, M. Malcangio, T. Malm, R. Mancuso, T. Masuda, M. Matteoli, B. W. McColl, V. E. Miron, A. V. Molofsky, M. Monje, E. Mracsko, A. Nadjar, J. J. Neher, U. Neniskyte, H. Neumann, M. Noda, B. Peng, F. Peri, V. H. Perry, P. G. Popovich, C. Pridans, J. Priller, M. Prinz, D. Ragozzino, R. M. Ransohoff, M. W. Salter, A. Schaefer, D. P. Schafer, M. Schwartz, M. Simons, C. J. Smith, W. J. Streit, T. L. Tay, L. H. Tsai, A. Verkhratsky, R. von Bernhardi, H. Wake, V. Wittamer, S. A. Wolf, L. J. Wu, and T. Wyss-Coray. 2022. ‘Microglia states and nomenclature: A field at its crossroads’, Neuron, 110: 3458–83.

35. Shinozaki, Y., K. Shibata, K. Yoshida, E. Shigetomi, C. Gachet, K. Ikenaka, K. F. Tanaka, and S. Koizumi. 2017. ‘Transformation of Astrocytes to a Neuroprotective Phenotype by Microglia via P2Y1 Receptor Downregulation’, Cell Rep, 19: 1151–64.

36. Sofroniew, M. V. 2015. ‘Astrocyte barriers to neurotoxic inflammation’, Nat Rev Neurosci, 16: 249–63.

37. Sozmen, E. G., J. D. Hinman, and S. T. Carmichael. 2012. ‘Models that matter: white matter stroke models’, Neurotherapeutics, 9: 349–58.

38. Suda, Y., T. Nakashima, H. Matsumoto, D. Sato, S. Nagano, H. Mikata, S. Yoshida, K. Tanaka, Y. Hamada, N. Kuzumaki, and M. Narita. 2021. ‘Normal aging induces PD-1-enriched exhausted microglia and A1-like reactive astrocytes in the hypothalamus’, Biochem Biophys Res Commun, 541: 22–29.

39. van Rooij, F. G., R. P. Kessels, E. Richard, F. E. De Leeuw, and E. J. van Dijk. 2016. ‘Cognitive Impairment in Transient Ischemic Attack Patients: A Systematic Review’, Cerebrovasc Dis, 42: 1–9.

40. Vermeer, S. E., W. T. Longstreth, Jr., and P. J. Koudstaal. 2007. ‘Silent brain infarcts: a systematic review’, Lancet Neurol, 6: 611–9.

41. Wang, Y., R. K. Leak, and G. Cao. 2022. ‘Microglia-mediated neuroinflammation and neuroplasticity after stroke’, Front Cell Neurosci, 16: 980722.

42. Witcher, K. G., C. E. Bray, J. E. Dziabis, D. B. McKim, B. N. Benner, R. K. Rowe, O. N. Kokiko-Cochran, P. G. Popovich, J. Lifshitz, D. S. Eiferman, and J. P. Godbout. 2018. ‘Traumatic brain injury-induced neuronal damage in the somatosensory cortex causes formation of rod-shaped microglia that promote astrogliosis and persistent neuroinflammation’, Glia, 66: 2719–36.

43. Xiong, X. Y., L. Liu, and Q. W. Yang. 2016. ‘Functions and mechanisms of microglia/macrophages in neuroinflammation and neurogenesis after stroke’, Prog Neurobiol, 142: 23–44.

44. Zhang, Q., C. Liu, R. Shi, S. Zhou, H. Shan, L. Deng, T. Chen, Y. Guo, Z. Zhang, G. Y. Yang, Y. Wang, and Y. Tang. 2022. ‘Blocking C3d(+)/GFAP(+) A1 Astrocyte Conversion with Semaglutide Attenuates Blood-Brain Barrier Disruption in Mice after Ischemic Stroke’, Aging Dis, 13: 943–59.

