## Supplemental Table 1 for "Mild subcortical stroke induces widespread astrogliosis independent of microglia and age"

| | | | Intensity ratio (I:C) | Cell counts (cells/mm2) | | N |
|---|---|---|---|---|---|---|
| | | | | Contralateral | Ipsilesional | |
| | | | Mean ± S.E.M. | Mean ± S.E.M. | Mean ± S.E.M. | |
| GFAP | 1 day | ACA | 1.01 ± 0.01 | 29.50 ± 22.70 | 24.97 ± 2.27 | 3 |
| | | MCA | 1.05 ± 0.01 | 70.36 ± 23.03 | 93.05 ± 28.44 | 3 |
| | | Striatum | 1.13 ± 0.03 | 93.05 ± 32.02 | 202.00 ± 37.36 | 3 |
| | 3 day | ACA | 1.06 ± 0.03 | 10.21 ± 4.40 | 212.78 ± 71.76 | 4 |
| | | MCA | 1.03 ± 0.03 | 51.07 ± 16.33 | 255.33 ± 30.00 | 4 |
| | | Striatum | 1.17 ± 0.05 | 76.60 ± 36.18 | 394.91 ± 71.41 | 4 |
| | 7 day | ACA | 1.04 ± 0.01 | 51.75 ± 11.91 | 133.45 ± 35.09 | 5 |
| | | MCA | 1.12 ± 0.04 | 84.43 ± 31.54 | 360.87 ± 107.16 | 5 |
| | | Striatum | 1.23 ± 0.05 | 119.15 ± 54.01 | 798.34 ± 140.24 | 5 |


| | | | | | | |
|---|---|---|---|---|---|---|
| GFAP | 1 day | ACA | 1.01 ± 0.01 | 29.50 ± 22.70 | 24.97 ± 2.27 | 3 |
| | | MCA | 1.05 ± 0.01 | 70.36 ± 23.03 | 93.05 ± 28.44 | 3 |
| | | Striatum | 1.13 ± 0.03 | 93.05 ± 32.02 | 202.00 ± 37.36 | 3 |
| | 3 day | ACA | 1.06 ± 0.03 | 10.21 ± 4.40 | 212.78 ± 71.76 | 4 |
| | | MCA | 1.03 ± 0.03 | 51.07 ± 16.33 | 255.33 ± 30.00 | 4 |
| | | Striatum | 1.17 ± 0.05 | 76.60 ± 36.18 | 394.91 ± 71.41 | 4 |
| | 7 day | ACA | 1.04 ± 0.01 | 51.75 ± 11.91 | 133.45 ± 35.09 | 5 |
| | | MCA | 1.12 ± 0.04 | 84.43 ± 31.54 | 360.87 ± 107.16 | 5 |
| | | Striatum | 1.23 ± 0.05 | 119.15 ± 54.01 | 798.34 ± 140.24 | 5 |


| | | | | | | |
|---|---|---|---|---|---|---|
| Iba1 | 1 day | ACA | 1.02 ± 0.01 | 86.36 ± 27.73 | 95.79 ± 33.06 | 4 |
| | | MCA | 1.01 ± 0.01 | 99.98 ± 36.46 | 181.98 ± 58.81 | 4 |
| | | border Striatum | 0.98 ± 0.01 | 107.70 ± 33.52 | 175.41 ± 60.33 | 4 |
| | | infarct (Striatum) | 0.96 ± 0.02 | | 140.55 ± 47.71 | 4 |
| | 3 day | ACA | 1.01 ± 0.00 | 80.00 ± 28.26 | 180.43 ± 88.71 | 4 |
| | | MCA | 1.04 ± 0.01 | 93.62 ± 12.24 | 200.86 ± 39.36 | 4 |
| | | border Striatum | 1.04 ± 0.02 | 107.24 ± 42.10 | 406.83 ± 61.30 | 4 |
| | | infarct (Striatum) | 1.02 ± 0.02 | | 461.30 ± 108.14 | 4 |
| | 7 day | ACA | 0.99 ± 0.01 | 98.05 ± 19.92 | 114.39 ± 20.13 | 5 |
| | | MCA | 1.01 ± 0.03 | 104.86 ± 28.04 | 385.38 ± 159.05 | 5 |
| | | border Striatum | 1.06 ± 0.03 | 114.39 ± 14.82 | 578.75 ± 254.44 | 5 |
| | | infarct (Striatum) | 1.19 ± 0.05 | | 1192.91 ± 89.05 | 5 |
