## Supplemental Table 2 for "Mild subcortical stroke induces widespread astrogliosis independent of microglia and age"

| | | Intensity ratio (I:C) | | | | Cell counts (cells/mm2) | | | | |
|---|---|---|---|---|---|---|---|---|---|---|
| | | MSCI | | sham | | Contralateral | Ipsilesional | | sham | |
| | | Mean ± S.E.M. | N | Mean ± S.E.M. | N | Mean ± S.E.M. | Mean ± S.E.M. | N | Mean ± S.E.M. | N |
| **GFAP** | **ACA** | 1.06 ± 0.02 | 6 | 1.00 ± 0.02 | 4 | 78.30 ± 22.91 | 174.76 ± 52.29 | 6 | 48.51 ± 3.77 | 4 |
| | **MCA** | 1.16 ± 0.02 | 5 | 1.02 ± 0.04 | 4 | 77.62 ± 7.01 | 399.00 ± 39.21 | 5 | 48.51 ± 15.50 | 4 |
| | **Striatum** | 1.21 ± 0.02 | 6 | 0.99 ± 0.01 | 4 | 290.51 ± 42.87 | 473.21 ± 31.04 | 6 | 252.78 ± 33.66 | 4 |
| **Vimentin** | **ACA** | 1.01 ± 0.01 | 6 | 1.00 ± 0.00 | 4 | 2.27 ± 2.27 | 4.54 ± 4.54 | 6 | 11.92 ± 7.55 | 4 |
| | **MCA** | 1.07 ± 0.03 | 6 | 1.00 ± 0.01 | 4 | 0.00 ± 0.00 | 127.10 ± 46.14 | 6 | 0.00 ± 0.00 | 4 |
| | **Striatum** | 1.20 ± 0.05 | 5 | 0.99 ± 0.02 | 4 | 13.62 ± 9.39 | 381.98 ± 61.96 | 5 | 11.06 ± 3.51 | 4 |
| **Iba1** | **ACA** | 1.00 ± 0.00 | 6 | 1.00 ± 0.00 | 4 | 5.67 ± 2.09 | 7.94 ± 4.45 | 6 | 4.26 ± 2.55 | 4 |
| | **MCA** | 1.00 ± 0.00 | 6 | 1.00 ± 0.00 | 4 | 1.13 ± 1.13 | 44.26 ± 26.59 | 6 | 2.55 ± 0.85 | 4 |
| | **Striatum** | 1.01 ± 0.01 | 5 | 1.01 ± 0.01 | 4 | 0.00 ± 0.00 | 66.73 ± 46.65 | 5 | 7.66 ± 2.55 | 4 |
