## Supplemental Table 3 for "Mild subcortical stroke induces widespread astrogliosis independent of microglia and age"

| | | Intensity ratio (I:C) | | | | Cell counts (cells/mm2) | | | | |
|---|---|---|---|---|---|---|---|---|---|---|
| | | MSCI | | sham | | Contralateral | Ipsilesional | | sham | |
| | | Mean ± S.E.M. | N | Mean ± S.E.M. | N | Mean ± S.E.M. | Mean ± S.E.M. | N | Mean ± S.E.M. | N |
| **GFAP** | **ACA** | 1.10 ± 0.01 | 3 | 1.02 ± 0.02 | 4 | 108.94 ± 68.20 | 211.07 ± 138.93 | 3 | 81.71 ± 25.17 | 4 |
| | **MCA** | 1.12 ± 0.01 | 3 | 1.03 ± 0.04 | 4 | 120.29 ± 57.28 | 267.81 ± 118.48 | 3 | 83.41 ± 26.46 | 4 |
| | **Striatum** | 1.13 ± 0.03 | 3 | 1.00 ± 0.05 | 4 | 403.99 ± 73.05 | 587.83 ± 92.14 | 3 | 280.44 ± 72.76 | 4 |
| **Vimentin** | **ACA** | 1.01 ± 0.02 | 3 | 1.01 ± 0.01 | 4 | 9.08 ± 6.00 | 65.82 ± 55.92 | 3 | 13.62 ± 5.01 | 4 |
| | **MCA** | 1.30 ± 0.22 | 3 | 1.00 ± 0.00 | 4 | 6.81 ± 3.93 | 240.58 ± 104.18 | 3 | 10.21 ± 9.11 | 4 |
| | **Striatum** | 1.87 ± 0.07 | 3 | 1.06 ± 0.04 | 4 | 111.21 ± 32.97 | 313.21 ± 14.17 | 3 | 51.07 ± 11.54 | 4 |
| **Iba1** | **ACA** | 1.03 ± 0.02 | 3 | 1.01 ± 0.01 | 4 | 108.94 ± 7.86 | 154.33 ± 59.53 | 3 | 142.99 ± 5.01 | 4 |
| | **MCA** | 1.08 ± 0.07 | 3 | 1.00 ± 0.01 | 4 | 165.68 ± 19.39 | 320.02 ± 188.49 | 3 | 159.16 ± 18.97 | 4 |
| | **border Striatum** | 1.16 ± 0.03 | 3 | 1.01 ± 0.00 | 4 | 167.95 ± 37.36 | 691.10 ± 154.38 | 3 | 191.92 ± 28.22 | 4 |
| | **infarct (Striatum)** | 1.34 ± 0.09 | 3 | | | | 1221.05 ± 181.74 | 3 | | |
